# Structural dynamics of mitochondrial ATP synthase in *Chlamydomonas reinhardtii* revealed by *in situ* CryoET

**DOI:** 10.1101/2025.09.10.674987

**Authors:** Emily Capper, Muyuan Chen

**Affiliations:** Division of CryoEM and Bioimaging, SSRL, SLAC National Accelerator Laboratory, Stanford University, Menlo Park, CA 94025, USA; College of Arts, Sciences, and Engineering, University of Rochester, Rochester, NY 14627, USA

## Abstract

The turbine dynamics of mitochondrial ATP synthases is an unresolved topic which *in situ* cryo-ET is poised to answer. Using publicly available tomogram data of vitrified *Chlamydomonas reinhardtii* cells, a monomer map of ATP synthase was refined to 6.67Å. With extensive heterogeneity analysis, we characterized the *in situ* conformation states of F1 head, central stalk, and upper peripheral stalk (UPS) individually, and analyzed the by-particle correlation between movement of the different domains, revealing the complex dynamics of the system within cells. While oligomycin sensitivity conferral protein (OSCP) and the UPS were found to bend with F1 head rotation, coupling between the rotation of central stalk and F1 head rotational states, suggested by previous *in vitro* studies, were not detected from ATP synthase inside mitochondria.

## Introduction

Mitochondria are the primary generators of ATP for essential cellular processes. Complexes I-IV, embedded in the inner mitochondrial membrane (IMM), use electrons imported on carrier molecules to increase the concentration of protons in the intermembrane space. ATP synthase (aka complex V) releases the proton gradient across the IMM to create rotational force. Such force liberates tightly-bound ATP after its synthesis from ADP and phosphate in the active site.

Here, considering F-ATPases ^1–3^, the specific mechanism converting proton motive force to ATP is not comprehensively understood, but many facts are clear: the *c*-ring, assembled from 8-17 *c*-subunits and embedded in the IMM, is the rotor that protons drive the rotation of before being released into the matrix ^1,4,5^. The *a*-subunit (*ATP6*) ^6^, with a large contact surface on the *c*-ring within the IMM ^5,7–9^, guides protons into the rotor and blocks their short-circuiting, driving productive rotation the long way around ^1,4,10^. The *c*-ring is connected to a central shaft matrix-side ^1,9^, responsible for driving catalysis asymmetrically in the F1 head ^5,7^, and is mechanistically reversed for V-ATPases ^11^. This pseudo-c3 hexamer consists of alternating alpha and beta subunits (three each), though beta is the main site of catalysis ^5,7,11–14^. To prevent unproductive spinning, a peripheral stalk serving as the stator is anchored into the membrane with a flexible linker attached to the top to retain the F1 head ^9,11,12^.

However, the proteins which compose the peripheral and central stalks can vary considerably between taxa: while bacterial species’ peripheral stalks have two matrix proteins ^1,5^, *Polytomella* sp. has seven ^7^—the latter species being closest to the subject of this work, *Chlamydomonas reinhardtii*, with a high-resolution structure of ATP synthase. In *C. reinhardtii*, the central stalk is composed of delta, epsilon, and gamma proteins ^15^, the latter of which reaches into the F1 head. Totalling findings from *C. reinhardtii* and *Polytomella* sp., the ASA1-10 proteins compose the peripheral stalk ^15–22^ (meeting the *a*-subunit in the membrane) and oligomycin sensitivity conferral protein (OSCP) is linked to the F1 head ^5^ by N-terminal domains of beta, one of which links back to the stator ^7^. The use of the ATP synthase-associated (ASA) proteins to compose the peripheral stalk in chlorophycean algae is unlike other eukaryotes, which do not possess them ^5,15,16,22^.

Though there are tetramer ^23^ and hexamer ^24,25^ arrangements of ATP synthase in more distant species, the mitochondrial ATP synthase of *C. reinhardtii*’s close relative *Polytomella* sp. forms dimers. Its dimers are arranged long-ways and “back-to-back,” with stators facing each other. They line the cristae in a helical series and provide for their curvature, since the dimer structure necessarily entails being in a bent part of the membrane ^1,8,14,15^.

Whereas single-particle cryo-EM is well-suited for producing 2D data of purified *in vitro* systems, cryo-ET allows rendering a volume model from cross-sectional electron micrograph images taken as the vitrified subject sample is tilted laterally. Because vitrification freezes proteins in-place, data from whole cells represents states and locations of proteins *in situ* ^8,26^. Dynamic proteins altogether exist in a distribution of states, so when their data is aligned to form a density map, regions with the most movement will necessarily be lower resolution. By mapping the span of heterogeneous states to a latent scale by-particle, models of intermediate states can be separately resolved.

Most work generating cryo-EM maps for mechanistic analysis of V- and F-ATPases use data taken from purified systems, with a more limited *in situ* literature. The former, which includes structures from *Polytomella* sp. ^7^, *S. cerevisiae* ^11^, “bovine” ^27^, and *G. stearothermophilus* ^28^, all show evidence of peripheral stalk bending (depending on how large the stalk is), F1 head rocking, and central stalk rotation. The *in vitro* argument for 30° central/stalk F1 head–coupled rotation in *Polytomella* sp. was made on the basis of observing the necessary rotational states and that OSCP binders affect the ATP synthase Michaelis constant, so (stopping) the movement of the F1 head is mechanistically relevant ^7^. These *in vitro* states represent possible mechanistic states, but the distributions are not necessarily representative of *in situ* dynamics. The only *in situ* literature concerning ATP synthase dynamics, in *Polytomella* sp., shows that there exist dynamic states of the central stalk, F1 head, and that F1 head states can be derived from within central stalk rotational classes ^8^. However, it does not itself make the strong claim made by the *in vitro* work ^7^. A critical demonstration of coupled movement would be a sinusoidal (or any) correlation of the F1 head rotation as a function of central stalk rotation.

In this work, we use cryo-electron tomograms of mitochondria in *C. reinhardtii* published by Thermo Fisher Eindhoven ^29^—and recently used to describe other proteins of the ETC ^26^—to subtomogram-average ATP synthase particles into an average 6.67Å monomer map. We also perform heterogeneity analysis to derive maps representing dynamic states and assign dynamic states by-particle, finding no evidence of F1 head–central stalk coupled rotation, instead showing OSCP–F1 head correlated movement.

## Results

We selected 93 of the mitochondria-focused tomograms of xenon plasma–FIB milled lamellae with vitrified whole cells published by Thermo Fisher Eindhoven ^29^. The chosen mitochondrial tomograms were then screened manually for a few hundred ATP synthase dimer particles, along the ridges of cristae, which were refined into an initial model. EMAN2 template matching selected several thousand more particles across all tomograms using the initial model, then refined into a higher-resolution dimer model.

Dimer model particles were re-extracted for monomers, then went through several stages of refinement including Gaussian mixture model (GMM) based orientation refinement ^30^ before reaching the final 6.67Å monomer map. The resolution of the monomer map was lower in more dynamic regions (Figure 1A, S1).

**Figure 1.**
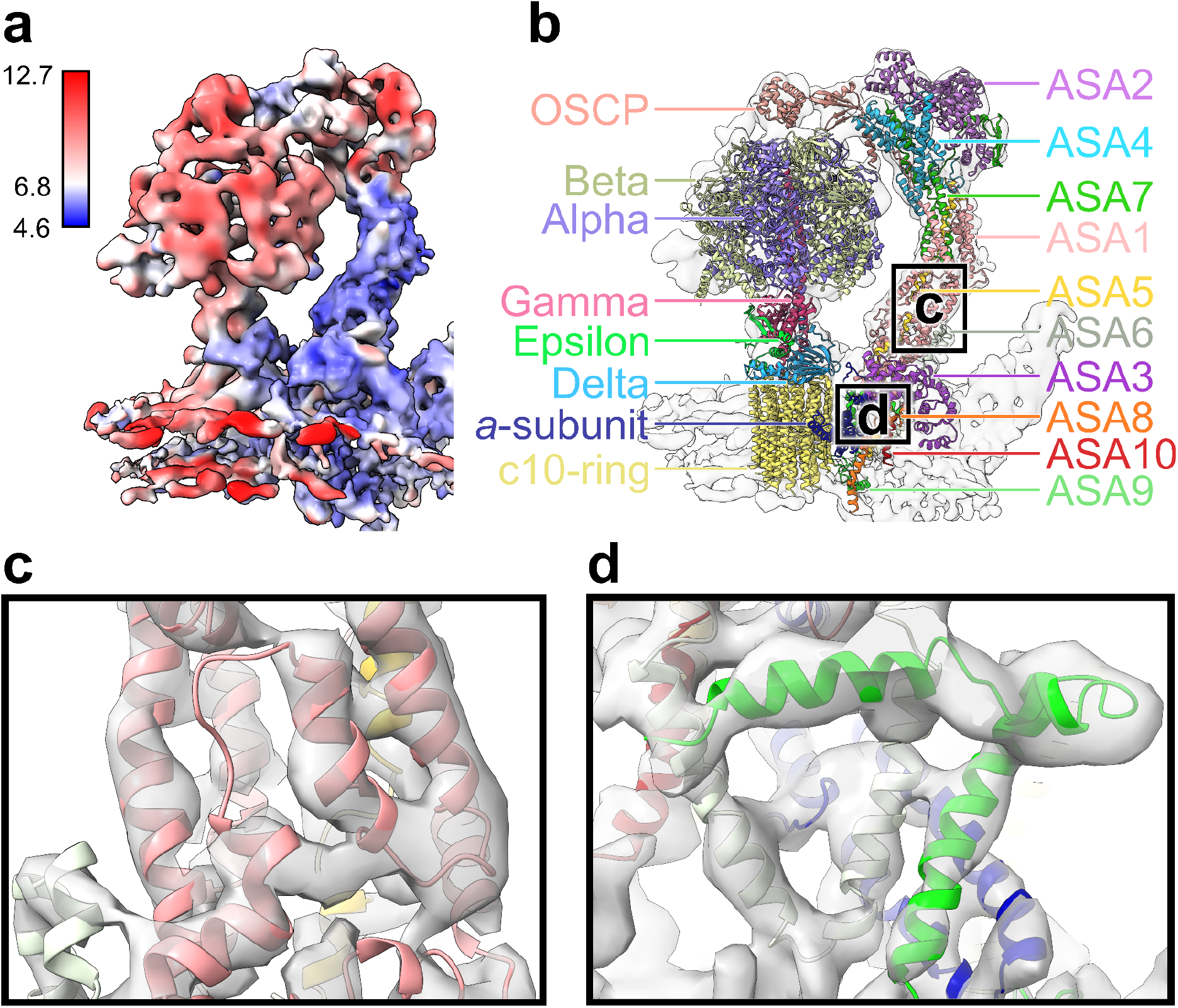
*Chlamydomonas reinhardtii* ATP synthase monomer model. (a) Map with surface colored according to its resolution in angstroms. Regions of highest resolution, in blue, correspond to the peripheral stalk and membrane proteins. Regions of lower resolution, in red and white, include the UPS, the F1 head, and the central stalk. (b) The ribbon version of the molecular model fitted to the density map is shown and labeled by-protein in multicolor. Overlap with the outline of the map is lower in regions of lower resolution. (c) First close-up of b, several alpha helices in the lower peripheral stalk. ASA1/5/6 are visible. (d) Second close-up of b, alpha helices of proteins in the membrane-embedded region. The *a*-subunit and ASA6/9/10 are visible.

With assistance of the ATP synthase molecular model from *Polytomella* sp., AI-folded ^31,32^ native *Chlamydomonas reinhardtii* proteins were fitted into the monomer map derived here (Figure 1B). In regions of high resolution, particularly the peripheral stalk and parts of the membrane-embedded complex, alpha helices of several homologous protein pairs have similarly strong correspondence to the map, among them ASA1/6 (Figure 1C), ASA6/9, and the *a*-subunit (Figure 1D). OCSP is mapped at lower resolution, but also shows strong correlation (Figure 2A).

**Figure 2.**
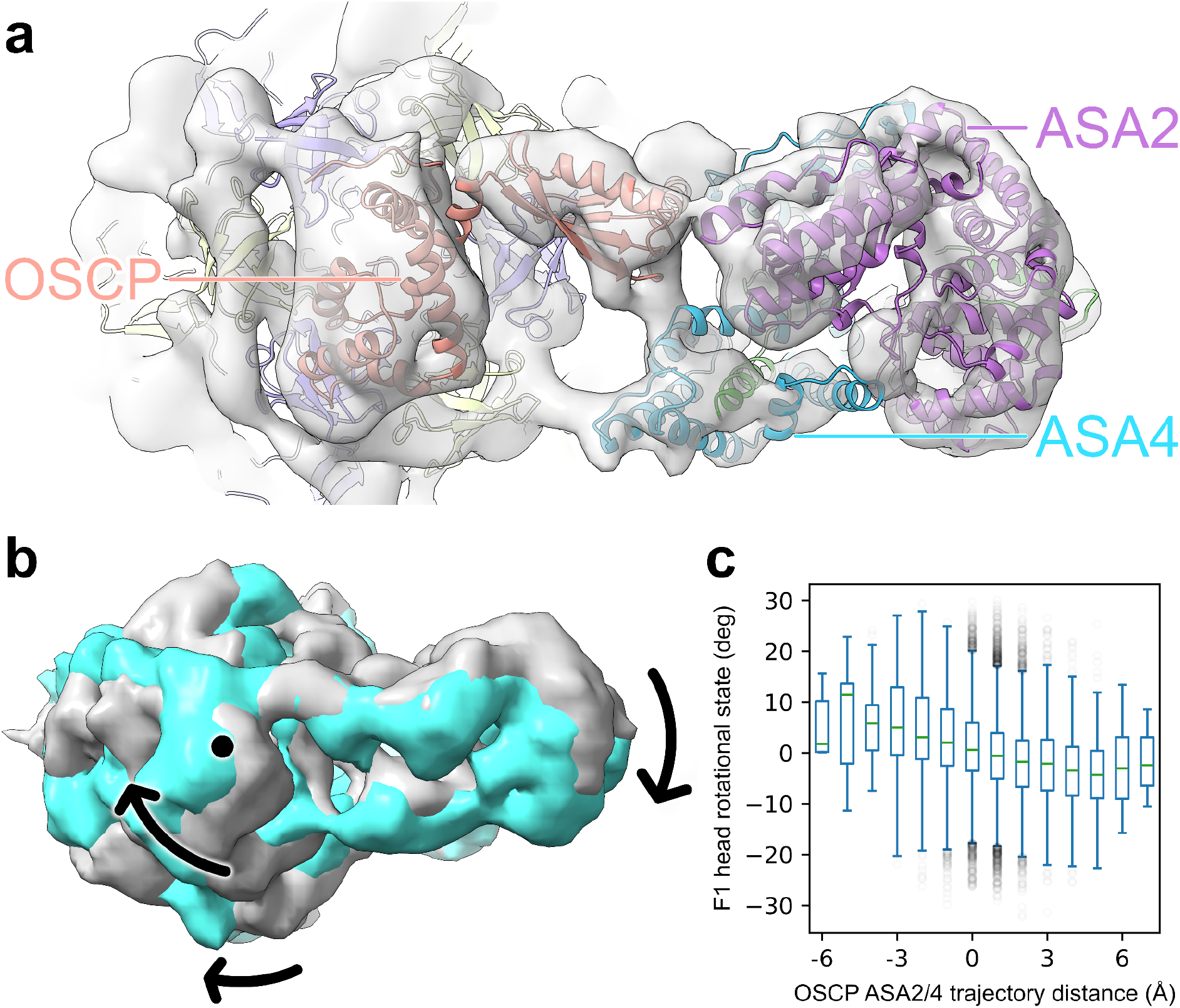
OSCP ASA2/4 movement trajectory corresponds to rotation of the F1 head. (a) Top view of UPS ribbon proteins fitted to the map, with labels for OSCP and ASA2/4, and in the same orientation as b. (b) Overlaid maps showing the largest difference in resolved positions of the F1 head and the UPS. Labeled with arrows, clockwise rotation of the F1 head about the central stalk axis, labeled with a dot, corresponds to clockwise movement of the UPS around the same. (c) Parallel box plot showing the relationship between 1Å buckets of OSCP ASA2/4 trajectory distance and their respective particles’ F1 head rotational state average. OSCP ASA2/4 movement was standardized to movement of the ASA2-Leu215 α-carbon. Outliers of fat-tailed distributions are shown, faded.

As previously anticipated ^33^, we find that *C. reinhardtii* ATP synthase lacks a “bridge” between monomers, present in *Polytomella* sp. ^7,8^. Though our model does not incorporate the linking of alpha subunits’ N-termini to OSCP or the upper peripheral stalk (UPS), the lower-resolution map densities (Figure 2A) imply it exists here too. Previous algal biochemical results have shown subunit interactions ^18–21^ which are confirmed by our and previous ^7^ molecular fitting. Subunit location, size, and accession is shown in Table S1.

In contrast to cryo-EM results of purified *S. cerevisiae* ATP synthase, where evidence was provided for vast elastic deformation of the peripheral stalk being associated with catalytic action generating ATP ^11^, stator movement observed in these data is not nearly as dramatic, as the alpha helices are clearly distinguished without any particle classification. The same applies to the *Polytomella* sp. structure ^7,8^, a more homologous point of comparison for *C. reinhardtii*.

### General results on ATP synthase movement

Previous *in vitro* and *in situ* studies analyzed the dynamics of the ATP synthase by grouping the particles into multiple discrete classes corresponding to the conformation of the complex. Despite its relevance to the dynamics hypothesis, the by-particle distributions of subunits’ orientations are not discussed in prior works. Here, using the GMM based heterogeneity analysis approach ^34^, each oriented particle is aligned about the central axis of the monomer model, and the by-particle angular deviation of the F1 head region is calculated. F1 head rotation does not occur in revolutions, because it is constrained by OSCP and the stator, so the range of its distribution is necessarily less than 360° and less than 90° from literature results of *Polytomella* sp. ^5,7,8^. From our calculation, the F1 head rotational distribution centers at 0° (set as the center angle owing to it being the resolved average rotational state), with 95% of particles being within ±19° (Figure 3C).

**Figure 3.**
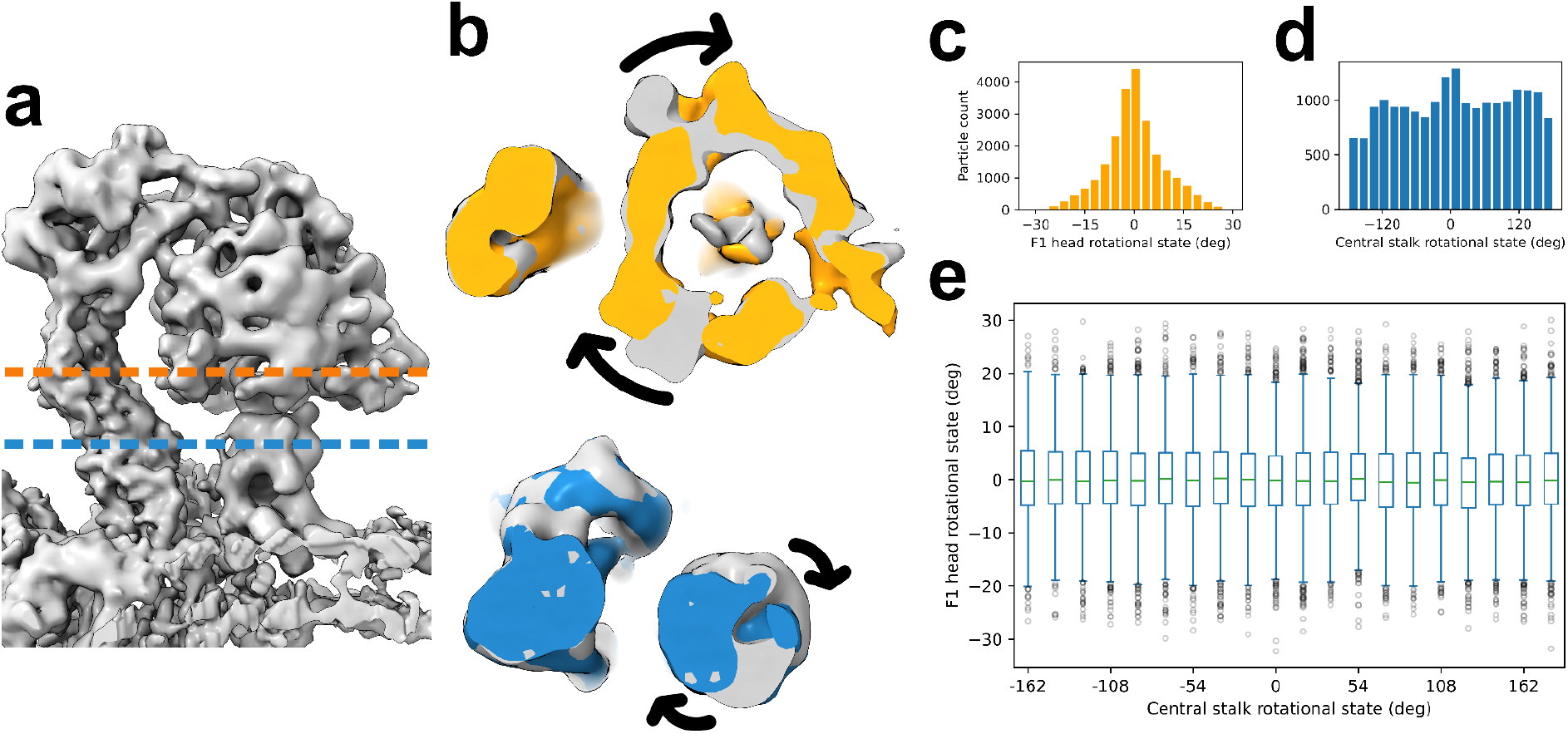
F1 head and central stalk rotation is uncorrelated. (a) Side view of the grey monomer map, with dashed lines indicating the planes of cross-section shown in b. (b) Cross-sections of maps showing rotation of the F1 head and central stalk, in orange and blue respectively, against another grey state. Arrows indicate direction of rotation between maps. (c) Distribution of F1 head rotational states by-particle, from the neutral position. (d) Distribution of central stalk rotational states by-particle. (e) No correlation between the F1 head and central stalk states is evident. KS tests comparing the F1 head distribution of each central stalk rotational group against the F1 head reference distribution in c showed no significance for any bucket (p=1).

Additionally, owing to its lack of rotational symmetry, the central stalk’s orientation is identifiable and classifiable in a similar manner. Orientations of the central stalk span across all 360 degrees, with a trimodal distribution due to pseudo-c3 catalytic action that several cryo-EM works on ATP synthase resolve states for ^7,8,11,12^—suggesting dwelling prior to higher-energy rotational states (with respect to the F1 head) for catalytic action. Within expectations, the central stalk angular distribution is multimodal and roughly evenly spaced, at 120°, 0°, and -120° (Figure 3D). In particular, the *Polytomella* sp. paper suggests one smaller (15°) and two larger (29°, 32°) primary rotary state dwells ^7^, which appear to be reflected here as one narrow and two wide rotational modes.

### Correlated OSCP–F1 head movement is observed

The OSCP subunit of ATP synthase is a flexible linker that connects the rotating F1 head domain to the stationary peripheral stalk, preventing unproductive “over-rotation” of F1. In previously reported structures, in both *Polytomella* sp. ^7^ and other eukaryotic species ^5,9,11,12^, OSCP serves as a flexible hinge. Considerable flexing is observed in species with only coiled-coil peripheral stalks ^28^, whereas in *Polytomella* sp. the bulky peripheral stalk is noted to stay rigid regardless of the movement of the F1 head, and only slightly flexes at the top ^7^. In our averaged structure, the upper peripheral stalk connecting to OSCP is not well resolved—including ASA2 and ASA4—suggesting movement of that domain, too.

By focusing the heterogeneity analysis on this region ^34^, we observe a continuous movement of the UPS, which rotates in coordination with the motion of the F1 head and the OSCP. The upper stator trajectory was reconstructed by selecting particles along the first eigenvector in the latent conformational space built by the deep neural networks, and each 3D map was reconstructed using a 1600-particle subset (Movie S1, Figure S2). From the stack of 3D maps, it was found that the ASA2/4 domain moves across 7Å clockwise (standardized to the ASA2-Leu215 α-carbon of the fitted molecular model) for 6.5° clockwise rotation of the F1 head (Figure 2B), with the middle 95% of particles having 5Å ASA2/4 movement for 6° F1 head rotation (Figure 2C).

The fact that the UPS is also flexible in the *in situ* mitochondria system (while staying rigid *vitro*) suggests this is a transient motion related to the dynamic of the system. It is hypothesized here that because OSCP controls the over-rotation of the F1 head, the excess energy of that over-rotation may be transmitted through the flexible hinge, and absorbed by the upper part of the peripheral stalk, leading to the observed flexibility.

### Self-correlation scores suggest c10 c-ring state

Existing evidence suggests a c10 *c*-ring configuration in *C. reinhardtii*: the c10 *Polytomella* sp. ATP synthase ^7^, older *C. reinhardtii* ATP synthase mass spectrometry data ^15^, and the identical membrane-embedded portions of the *c*-subunit between the species (Table S1). To further confirm this, as *c*-subunit count is relevant to central stalk angular binning, we use two indirect metrics. First, the *c*-ring map was reconstructed with each symmetry from c1 to c15 and then superimposed on the original for similarity score. A peak at c10 against decreasing scores implies a c10 configuration (Figure S3A). Next, the similarity of the unmodified *c*-ring volume was calculated against itself for every degree of rotation. 10 evenly-spaced similarity minima were observed (Figure S3B), also supporting c10.

### Correlated central stalk–F1 head motion is not observed

Recent single-particle cryo-EM models of purified ATP synthase from bovine ^27^ and *Polytomella* sp. ^7^ suggest that coupling between the central stalk and the F1 head occurs during catalysis, whereby they rotate together for 20-30°. Among algae—with bulky peripheral stalks—this would be compensated by OSCP and UPS bending. In mitochondrial tomograms of whole vitrified cells, however, we do not find clear and compelling evidence for that coupling hypothesis.

By a simple version of coupled rotation, one would expect a triperiodic correlation between their rotational states, fitting to *a* * *sin* (3*x* − *b*), for instance. The aligned and oriented ATP synthase monomer particles (Figure 3A) were grouped according to their F1 head (Figure 3B, 3C) and central stalk rotational states (Figure 3B, 3D), and the former was plotted against 20 bins each spanning 18° of central stalk rotational states. Unlike what was reported in the *in vitro* structures, no periodic correlation was observed (Figure 3E) and KS tests of each angular bucket against the F1 head reference distribution showed no significant difference (p=1), indicating a lack of stalk-head coupled rotation.

Whereas spatial resolution is a property in 3D-space for distinguishing meaningful features in the map, angular resolution of by-particle assignment is estimated here to mark the limit of higher-frequency correlations we would be able to observe. The maximum radius at which structural features exist from the central stalk axis is ∼30Å, so the central angle corresponding to the local resolution chord of ∼10Å is ∼19°, the angular resolution. Therefore, if 20-30° F1 head–central stalk coupling was occurring, then we would have been able to observe it, which we do not. However, this leaves open the possibility of coupling dynamics at higher rotation frequency that we cannot assess here.

Two primary hypotheses emerge to explain this observation: first, supposing this result is consistent with previous works’ findings of large F1 head–central stalk coupling. Like most ATP synthases ^1^, the 10 *c*-subunits in the *c*-ring and the three αβ catalytic subunits of the F1 head have a symmetry mismatch (pseudo-c10 and pseudo-c3). Alpha helical densities were distinguishable in the model’s *c*-ring, so its “discrete” motion is implied. Therefore, the c10-ring may rotate for three or four subunits to progress the catalytic action of the F1 head by one step (10/3 *c*-rotations being required for one catalytic subunit action).

If asymmetry in catalysis can occur at any of the three αβ subunits over the course of many catalytic cycles (example stages: store 2/3 *c*, use 1/3 *c*, use 1/3 *c*), and its timing across ATP synthase complexes is not necessarily in sync (the central stalk angle may vary by multiples of 120°), then for each case, alternate states of the F1 head should exist for the same central stalk state (and vice versa), and the flexible bending of the stator observed in more distant species—apparently to smooth over the symmetry mismatch and to store energy for the catalytic power stroke ^1,5,7,11^—would also apply here in principle. However, such a higher-order correlation, in addition to a subtler coupling, is beyond the resolution of the data here.

The alternate hypothesis suggests instead that the observed OSCP–F1 head coupling is the result of OSCP moving to account for mostly random (or otherwise central stalk–independent) F1 head motion, rather than variable resistance due to catalytic action. To account for the *c*-ring/F1 head symmetry mismatch, another stabilizing interaction must account for the over-rotation. For example, large and low-energy rotational dwellings in which there is a buffer for ≥2/3 *c* of rotation would eliminate the need for asymmetry-mediated F1 head–central stalk coupling.

Too, the models using cryo-EM images of purified, *in vitro* ATP synthase ^7,27^ can be reasonably construed as consistent with this hypothesis. Whereas tomograms of mitochondria in vitrified live cells tend to contain a distribution of ATP synthase states representative of its *in vivo* dynamics, the purified ATP synthase samples are more likely to inflate representation of low free energy states. This may include rocking of the F1 head and central stalk together within 15-32° energetic dwelling arcs at the beginning of each ∼120° catalytic stage, using OSCP to hinge ^7^. A wide dwelling arc would permit rotation, to an extent, of the F1 head (and trigger subsequent hinging of OSCP), without the central stalk needing to move concurrently.

### Correlated cross-dimer movement is not observed

The last finding establishes that ATP synthase dimers need only be considered as monomers for the purposes of their mechanics. Considering a dataset of ATP synthase dimer particles and assigning rotational states of F1 heads and central stalks for each monomer, rotational states of either of the same may be considered correlated within a dimer if, for a slice of particle data around one monomer’s rotational modes, the distribution of rotational states of the other monomer for the same particles is statistically indistinguishable from the reference distribution.

Comparisons were made between both F1 heads within a dimer (Figure S4A), but no statistically significant difference (KS test, p>0.4) between negative and positive angle groups was found; and between central stalks within a dimer (Figure S4B), with no significant difference (KS tests, p>0.2) between any pairs of particles within 30° of the three peak angular stalk rotational frequencies.

The only *in situ* cryo-ET of ATP synthase in *Polytomella* sp., using fewer discrete rotary states, also found no intra-dimer rotary state correlations, but did so by mapping those rotary state models back into a mitochondrial tomogram and visualizing ^8^. Though it may appear visually obvious from a single remapping, our statistical testing shows a lack of correlation across a larger set than can be found in one tomogram.

## Conclusions

Using a publicly available cryo-ET dataset of plunge-frozen live *Chlamydomonas reinhardtii* ^29^, in this study we apply heterogeneity analysis to several domains of mitochondrial ATP synthase, revealing the energetically representative states captured by rapid freezing. This property is not necessarily ensured in cryo-EM of *in vitro*, purified systems, but especially for the proton-gradient-dependent ATP synthase its *in vitro* states cannot reasonably be construed as representative of its mechanics, even when reconstituted in a membrane.

We demonstrate by-particle assignment of dynamic states along latent and rotational axes with deep learning–based structural heterogeneity algorithms ^30,34^. Rather than only presenting a series of discrete conformational states like in prior works ^7,8^, which is necessary but not sufficient to evidence a dynamics hypothesis, the fine-combed distributional nature of the data supporting our findings accurately represent the actual frequencies of possible conformational states, and enables the statistical analysis of coordinated movements within the same particle.

Furthermore, this work is a case in point of sharing and reprocessing cellular cryo-ET data being critical. Unlike datasets of purified single proteins, cellular tomograms possess a span of native environment-interacting protein dynamics likely to go beyond the scope of the original publication, reusable to make new discoveries.

## Methods

### Model refinement from tomograms

Of the publicly available tilt series published by Thermo Fisher Eindhoven (EMPIAR-11830) ^29^, 93 of those containing mitochondria were selected for this work. All tilt series in the public dataset were aligned and reconstructed in EMAN2 ^35^, and the ice thickness of each tomogram is estimated based on contrast distribution along the z axis. The tomograms were sorted by the ice thickness, and 93 thinnest tomograms (<150 nm) with visually identifiable mitochondrial features were selected. From the tomograms, a few hundred particles of ATP synthase dimers were manually selected by following the edge of cristae. These particles generated a primitive model, which was used to template match 17k more dimer particles. An initial model—showing three pairs of ATP synthase along a crista—was created from template matched particles, then filtered for duplicates to further refine a dimer-centered model. Monomer particles were re-extracted from dimer particles for monomer model refinement, then GMM refinement to increase resolution to 6.67Å. This process is summarized in Figure S5.

### Heterogeneity analysis of ATP synthase

Heterogeneity analysis of the ATP synthase monomer particles was performed in EMAN2, focusing on three individual parts separately. For the junction between OSCP and F1 head, the GMM and deep neural network (DNN)–based methods were used to map the particles to a 2D conformational latent space ^34^. A soft mask was used to focus the analysis on the target region, including the F1 head and the upper parts of the stalk. The series of reconstructions were obtained by sampling particles along the first eigenvector in the latent space, and each frame of the 3D movies (Movie S1-S3) were reconstructed from 1600 particles.

For the rotation of F1 head and the central stalk, since we expect rotation of the entire domain according to previous literatures, we use a variant of the GMM-DNN-based heterogeneity analysis method that uses the prior of rigid body motion of individual domains (described in ^36^). From the results of the analysis, we computed the rotation angle of the two domains around the z axis for each particle, and grouped particles by the rotation angles to generate the 3D movies representing the rotation of each piece. Additionally, using the method described in ^30^, the rotation angles estimated from each particle were converted to Euler angle assignment of the particle, which was used to improve the resolution of the averaged structure at the corresponding local regions.

### Molecular model procedure

Molecular structures of all subunits were first folded by AlphaFold2 ^31,32^, then rigidly fit into the final GMM-refined map using ChimeraX ^37^. Using the molecular dynamics simulation plugin ISOLDE ^38^, constraints were placed on secondary structures of AlphaFold-folded proteins, then simulated—with manual interference—to fit into densities of the map in regions of high resolution. In some locations, constraints were released to resemble the *Polytomella* sp. ATP synthase molecular structure ^7^, and proteins were trimmed at the ends in regions where the structure was low-confidence, lacked a homologous structure, and where the map was sufficiently low resolution. The final model is refined in EMAN2 ^39^ to improve the overall geometry. Native tools in ChimeraX were used to generate figures ^37^.

### Statistical tests

Statistical testing was performed in a Jupyter notebook using the KS test function from the SciPy package ^40^, on data organized with Pandas dataframes. For KS tests comparing by-particle central stalk angular distribution bins of F1 head angles to the F1 head reference distribution (n=22630), from left to right Figure 3E, the sample sizes were n=654, 650, 939, 1003, 939, 939, 897, 840, 983, 1209, 1288, 971, 928, 976, 971, 985, 1094, 1092, 1071, and 833. For intradimer rotational state KS testing, F1 head positive and negative angle groups had n=4582 and 4806 respectively, and sizes of n=1101, 1361, and 1256 for the -120°, 0°, and 120° central stalk rotational modes respectively.

## Acknowledgement

This research is supported by NIH grant R01GM150905.

## Supplementary materials

**Table S1.**
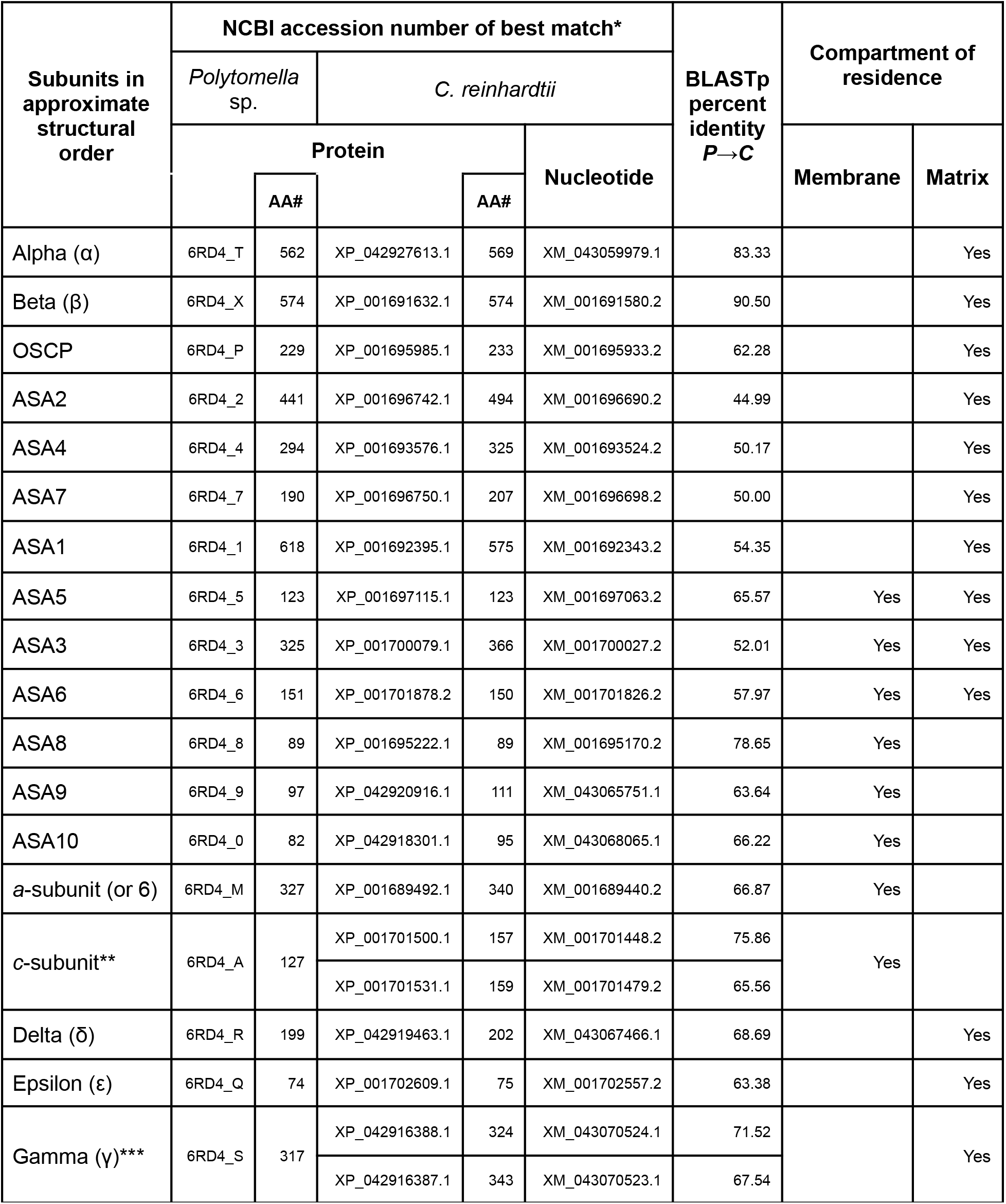
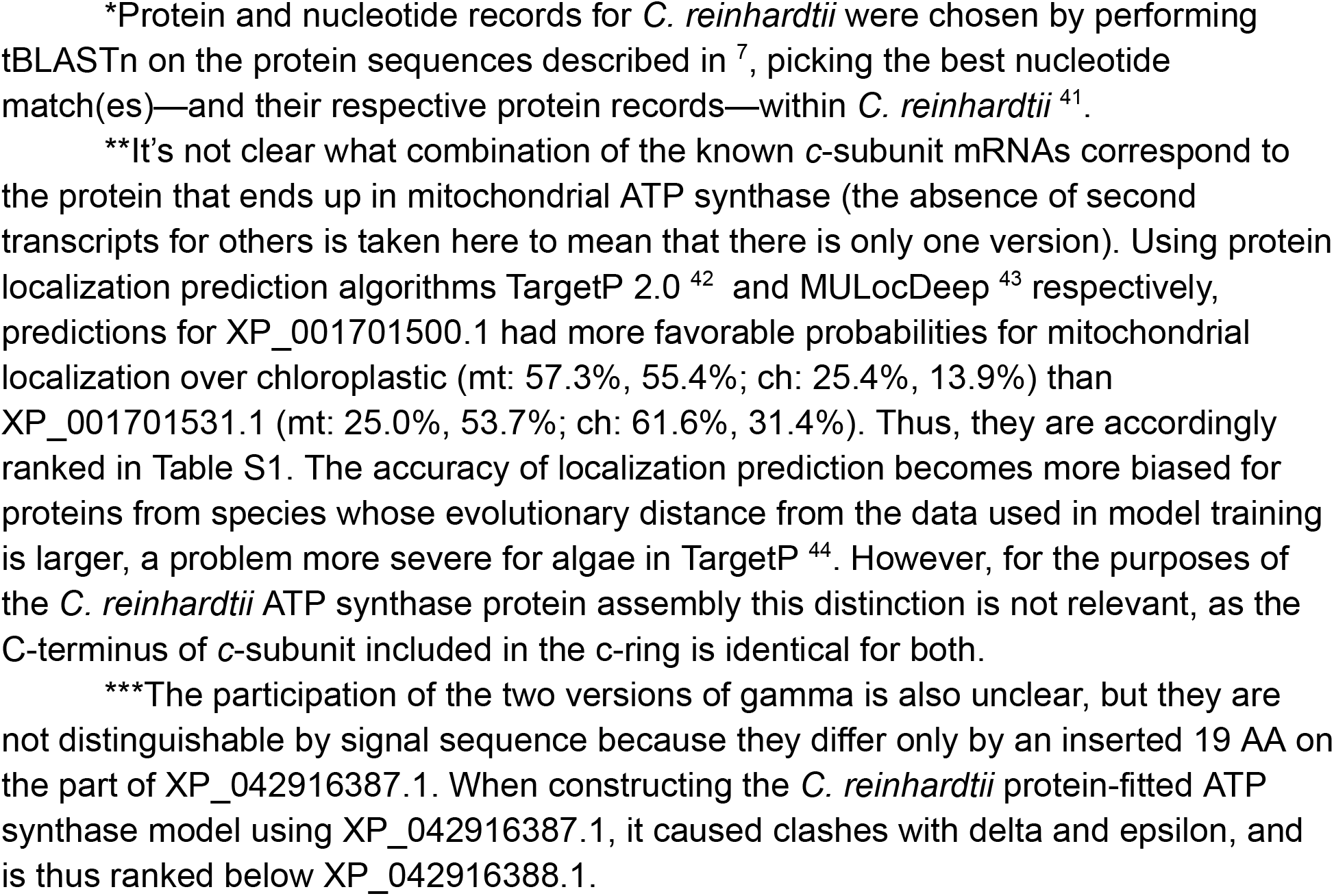
ATP synthase protein components of Chlorophycean algae.

| Subunits in approximate structural order | NCBI accession number of best match* | | | | | BLASTp percent identity $P \rightarrow C$ | Compartment of residence | |
| --- | --- | --- | --- | --- | --- | --- | --- | --- |
|  | *Polytomella* sp. |  | *C. reinhardtii* |  |  |  | Membrane | Matrix |
|  | Protein |  |  |  | Nucleotide |  |  |  |
|  |  | AA# |  | AA# |  |  |  |  |
| Alpha (α) | 6RD4_T | 562 | XP_042927613.1 | 569 | XM_043059979.1 | 83.33 |  | Yes |
| Beta (β) | 6RD4_X | 574 | XP_001691632.1 | 574 | XM_001691580.2 | 90.50 |  | Yes |
| OSCP | 6RD4_P | 229 | XP_001695985.1 | 233 | XM_001695933.2 | 62.28 |  | Yes |
| ASA2 | 6RD4_2 | 441 | XP_001696742.1 | 494 | XM_001696690.2 | 44.99 |  | Yes |
| ASA4 | 6RD4_4 | 294 | XP_001693576.1 | 325 | XM_001693524.2 | 50.17 |  | Yes |
| ASA7 | 6RD4_7 | 190 | XP_001696750.1 | 207 | XM_001696698.2 | 50.00 |  | Yes |
| ASA1 | 6RD4_1 | 618 | XP_001692395.1 | 575 | XM_001692343.2 | 54.35 |  | Yes |
| ASA5 | 6RD4_5 | 123 | XP_001697115.1 | 123 | XM_001697063.2 | 65.57 | Yes | Yes |
| ASA3 | 6RD4_3 | 325 | XP_001700079.1 | 366 | XM_001700027.2 | 52.01 | Yes | Yes |
| ASA6 | 6RD4_6 | 151 | XP_001701878.2 | 150 | XM_001701826.2 | 57.97 | Yes | Yes |
| ASA8 | 6RD4_8 | 89 | XP_001695222.1 | 89 | XM_001695170.2 | 78.65 | Yes |  |
| ASA9 | 6RD4_9 | 97 | XP_042920916.1 | 111 | XM_043065751.1 | 63.64 | Yes |  |
| ASA10 | 6RD4_0 | 82 | XP_042918301.1 | 95 | XM_043068065.1 | 66.22 | Yes |  |
| a-subunit (or 6) | 6RD4_M | 327 | XP_001689492.1 | 340 | XM_001689440.2 | 66.87 | Yes |  |
| c-subunit** | 6RD4_A | 127 | XP_001701500.1 | 157 | XM_001701448.2 | 75.86 | Yes |  |
|  |  |  | XP_001701531.1 | 159 | XM_001701479.2 | 65.56 |  |  |
| Delta (δ) | 6RD4_R | 199 | XP_042919463.1 | 202 | XM_043067466.1 | 68.69 |  | Yes |
| Epsilon (ε) | 6RD4_Q | 74 | XP_001702609.1 | 75 | XM_001702557.2 | 63.38 |  | Yes |
| Gamma (γ)*** | 6RD4_S | 317 | XP_042916388.1 | 324 | XM_043070524.1 | 71.52 |  | Yes |
|  |  |  | XP_042916387.1 | 343 | XM_043070523.1 | 67.54 |  |  |

\*Protein and nucleotide records for *C. reinhardtii* were chosen by performing tBLASTn on the protein sequences described in <sup>7</sup>, picking the best nucleotide match(es)—and their respective protein records—within *C. reinhardtii* <sup>41</sup>.
\*\*It's not clear what combination of the known *c*-subunit mRNAs correspond to the protein that ends up in mitochondrial ATP synthase (the absence of second transcripts for others is taken here to mean that there is only one version). Using protein localization prediction algorithms TargetP 2.0 <sup>42</sup> and MULocDeep <sup>43</sup> respectively, predictions for XP\_001701500.1 had more favorable probabilities for mitochondrial localization over chloroplastic (mt: 57.3%, 55.4%; ch: 25.4%, 13.9%) than XP\_001701531.1 (mt: 25.0%, 53.7%; ch: 61.6%, 31.4%). Thus, they are accordingly ranked in Table S1. The accuracy of localization prediction becomes more biased for proteins from species whose evolutionary distance from the data used in model training is larger, a problem more severe for algae in TargetP <sup>44</sup>. However, for the purposes of the *C. reinhardtii* ATP synthase protein assembly this distinction is not relevant, as the C-terminus of *c*-subunit included in the *c*-ring is identical for both.
\*\*\*The participation of the two versions of gamma is also unclear, but they are not distinguishable by signal sequence because they differ only by an inserted 19 AA on the part of XP\_042916387.1. When constructing the *C. reinhardtii* protein-fitted ATP synthase model using XP\_042916387.1, it caused clashes with delta and epsilon, and is thus ranked below XP\_042916388.1.

**Figure S1.**
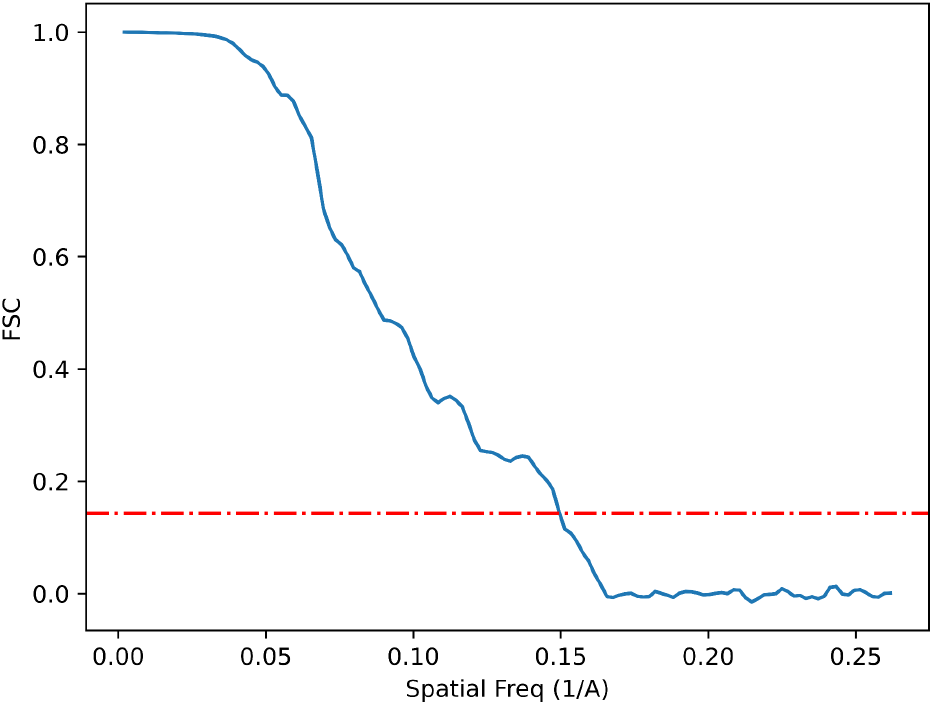
FSC resolution curve. At an FSC of 0.143, the resolution of the map in Figure 1 is 6.67Å.

**Figure S2.**
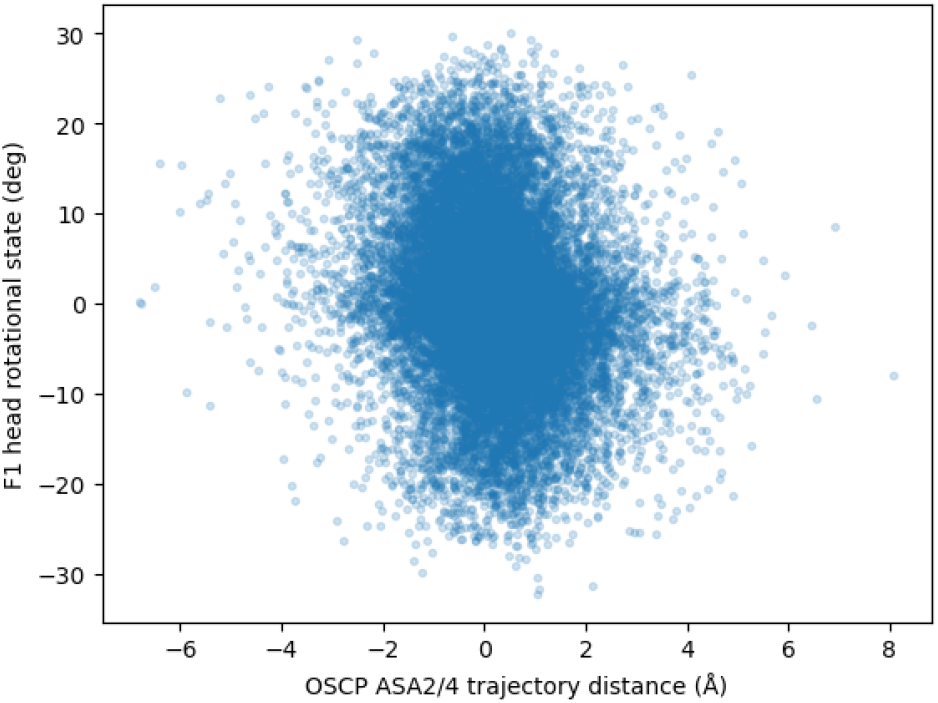
Particles’ F1 head rotational state against their DNN-assigned OSCP trajectory distance. A non-random relationship between the upper peripheral stalk trajectory distance and F1 head rotational state is shown, indicating the existence of correlated movement.

**Figure S3.**
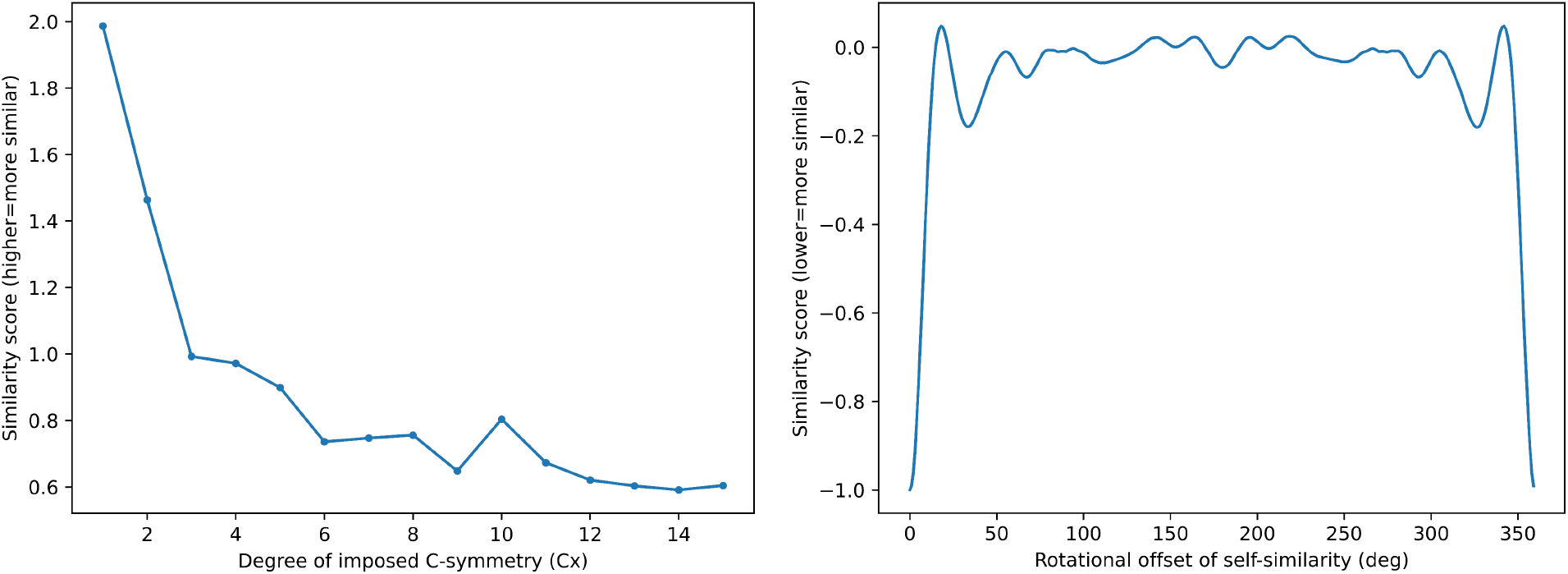
Similarity scores for imposed symmetry and rotational similarity performed on the *c*-ring volume. (a) Several C symmetries were imposed on the model’s *c*-ring and similarity to the original was calculated, where a higher score indicates more similarity. Note the peak at C10. (b) The *c*-ring volume was self-imposed and a similarity score calculated for each degree of rotation around the C symmetry axis, a lower score indicating more similarity. Note the 10 score minima.

**Figure S4.**
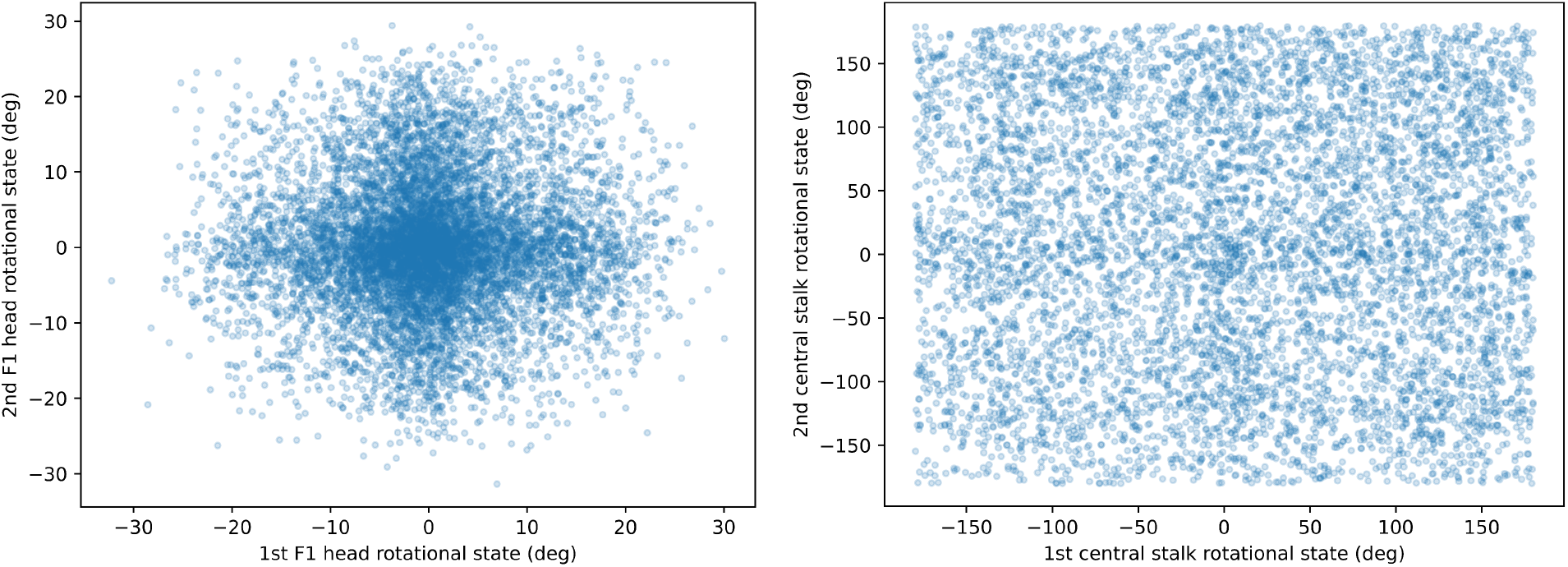
Correlations of intra-dimer F1 heads and central stalks. Each point represents a dimer particle with (a) F1 head and (b) central stalk rotational states assigned for each monomer.

**Figure S5.**
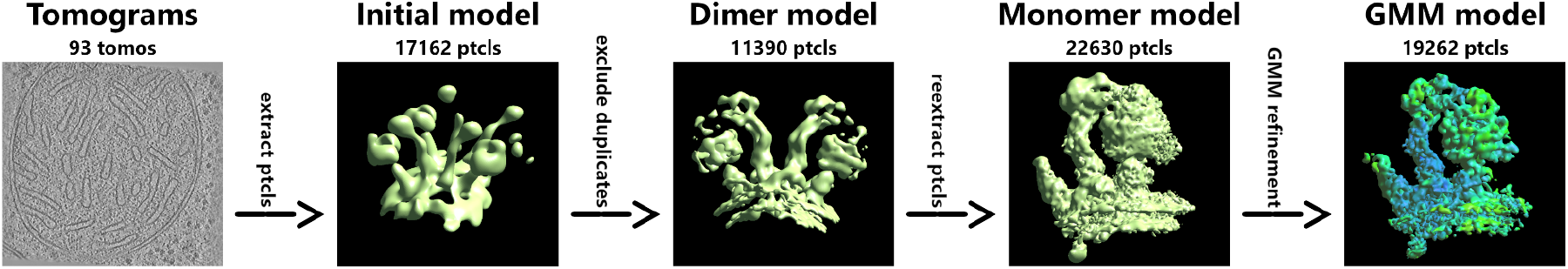
Subtomogram refinement process workflow.

